# The target landscape of N4-hydroxycytidine based on its chemical neighborhood

**DOI:** 10.1101/2020.03.30.016485

**Authors:** Jordi Mestres

## Abstract

N4-hydroxycytidine (NHC) has been recently reported to have promising antiviral activity against SARS-CoV-2. To join worldwide efforts in identifying potential drug targets against this pandemic, the target landscape of NHC was defined by extracting all known targets of its chemical neighborhood, including drugs, analogues, and metabolites, and by performing target predictions from two independent platforms, following the recent Public Health Assessment via Structural Evaluation (PHASE) protocol. The analysis provides a list of over 30 protein targets that could be useful in future design activities of new COVID-19 antivirals. The relevance for existing drugs within the same chemical space, such as remdesivir, is also discussed.

## Introduction

Scientists all over the world are joining efforts to fight the present SARS-CoV-2 pandemic, also referred to as Coronavirus disease 2019 or Covid-19. In a recent article on March 19, 2020, a prodrug of N4-hydroxycytidine (NHC; EIDD-1931) is shown to have antiviral activity against SARS-CoV-2 but also against multiple other endemic, epidemic and bat coronavirus [1]. In fact, the broad antiviral activity of NHC had been already explored widely in the past few months. On November 26, 2019, it was reported that NHC was able to inhibit murine hepatitis virus (MHV) and Middle East respiratory syndrome coronavirus (MERS-CoV) with minimal cytotoxicity [2]. A month earlier, on October 23, 2019, the same NHC prodrug (EIDD-2801) was found to be effective against multiple influenza virus strains [3]. And just over two years ago, on Jan 17, 2018, NHC was identified as a potent inhibitor of the Venezuelan equine encephalitis virus (VEEV) [4].

Results from those previous works associate the broad-spectrum antiviral activity of NHC and its prodrug (Figure 1) with increased transition mutation frequency in viral but not host cell RNA, supporting a mechanism of lethal mutagenesis [5]. It is accepted that nucleoside analogues increase the mutation rate of viral populations to levels incompatible with their survival [6]. However, how this lethal mutagenesis is achieved by NHC remains to be fully understood and multiple factors may play a role in it. In this respect, with worldwide efforts to identify potential targets against Covid-19 [7], exploring the target landscape of the surrounding chemical space of NHC may provide additional information on its mechanism of action that could guide the design of future lethal mutagenic agents.

**Figure 1.**
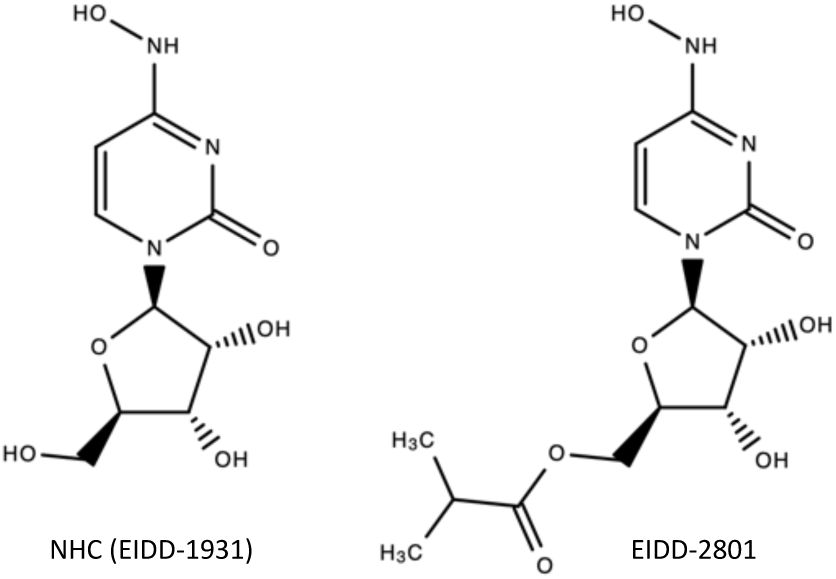
Structures of NHC (EIDD-1931) and its prodrug, EIDD-2801.

NHC was already identified as a deaminase inhibitor over 50 years ago [8]. Since then, a large amount of affinity data between small molecules and proteins has been accumulated into databases [9]. Under the similarity-property principle [10], the likely targets of any given molecule should be in consonance with the targets of its chemical neighborhood. This is partly the basis of recent initiatives such as the Public Health Assessment via Structural Evaluation (PHASE) developed by the US Food and Drug Administration’s Center for Drug Evaluation and Research (CDER) [11,12]. Following on that initiative, several similarity-based approaches will be used here to identify a candidate list of protein targets for NHC.

## Methodology

### Structural similarity

All similarities between pairs of molecules were computed using three types of two-dimensional descriptors implemented in CLARITY [13], namely, PHRAG, FPD and SHED, designed to describe chemical structures with different degrees of fuzziness [14,15]. Pharmacophoric fragments (PHRAG) are all possible fixed-length segments of five atom-features that can be extracted from the topology of a molecule [14]. In contrast, feature-pair distributions (FPD) capture the overall spreading of pairs of atom-centered features at different predefined bond lengths [14]. Finally, Shannon entropy descriptors (SHED) are derived from simplified FPD, in which, instead of using the actual feature-pair counts at each path length, the variability within all possible feature-pair distributions is quantified using the concept of Shannon entropy [15]. When using PHRAG and FPD, the similarity between two molecules corresponds to the overlapping fraction of their respective profiles [14], whereas with SHED, Euclidean distances are calculated instead [15]. The final similarity is obtained by computing the geometrical average between the two highest similarities from the three descriptors. The chemical neighborhood of any given molecule is defined by all molecules found within a similarity cut-off of 0.7.

### Metabolite prediction

Metabolites for NHC were predicted using the knowledge-based approach implemented in CLARITY [13]. The method relies on the information extracted from a set of 8,961 metabolic transformations collected for 1,791 drugs [16]. An all atom-by-atom superposition of all pairs of drug-metabolite structures allows for identifying the site of metabolism (SoM) and the type of chemical transformation produced, with details on the parts of the molecules that were added, deleted or modified. Then, we use PHRAG descriptors [14] to describe the atom environment surrounding the SoM in drugs associated with a given chemical transformation. Finally, a statistical analysis is performed to assess how often certain PHRAG descriptors are present in the SoM of each chemical transformation. PHRAG descriptor signatures could be defined for a total of 71 chemical transformations [16]. Their presence in chemical structures is transformed into a probability that the molecule would actually undergo a given chemical transformation and a confidence score is ultimately assigned to each metabolite predicted.

### Target prediction

Following on the framework defined recently by the PHASE initiative from CDER [11,12], targets for NHC were predicted using two platforms, namely, CLARITY [13] and SEA [17]. Both platforms use the two-dimensional structure of molecules to predict potential binding targets and their predictive performance was previously assessed by their respective developers [18,19]. SEA uses descriptor-based similarity to compare the structure of a molecule to the chemical structures with known *in vitro* binding affinity in ChEMBL [17]. Approximately 2,300 protein targets are covered. For each predicted target, a p-value and the similarity of the closest molecule are provided. CLARITY uses six ligand-based approaches that rely on descriptor-based molecular similarity (*vide supra*), an implementation of the similarity ensemble approach [17], fuzzy fragment-based mapping, quantitative structure-activity relationships, machine learning methods (including support vector machine, random forest and neural networks) and target cross-pharmacology indices [14]. The training set for the 4,799 protein target models is generated from *in vitro* affinity data contained in both public and patent sources [20]. For each target prediction, the projected affinity and mode of action are provided alongside with a confidence score based on the number and type of the methods that independently contribute to the prediction.

## Results & Discussion

### Drug neighborhood

A total of 14 chemical neighbors were identified, 8 sharing the same scaffold of NHC [21] and 6 having a different isosteric scaffold. Among those, there were three drugs, namely, cytarabine, azacytidine and adenosine (Figure 2). Since some of the proteins interacting with those drugs are already known, an analysis of their pharmacologies will help defining the target landscape around NHC.

**Figure 2.**
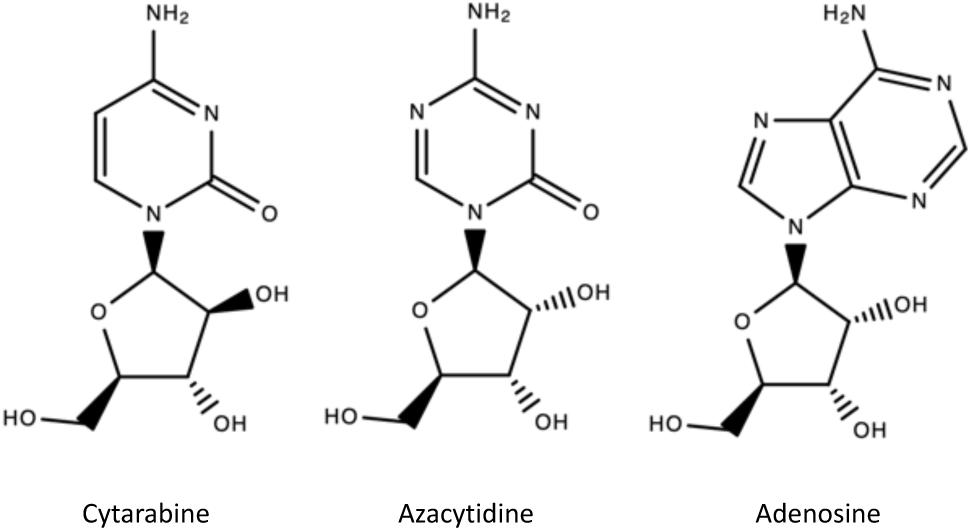
Structures of the three drugs in the chemical neighborhood of NHC.

Accordingly, all known drug-target interactions, in both public and patent databases available in CLARITY, were extracted and compiled in Table 1. As can be observed, there is almost no overlap in their known interacting proteins. This should not be too surprising if one considers that those molecules were most likely not tested consistently against the same panel of proteins [22]. Among the few known common targets, all three drugs bind to members of the adenosine receptor family and two of them, cytarabine and adenosine, are known to be active against the transmembrane domain-containing protein TMIGD3.

**Table 1.** List of known targets for the drugs in NHC’s chemical neighborhood.

| Protein / Protein family name | Gene(s) | Cytarabine | Azacitidine | Adenosine |
| --- | --- | --- | --- | --- |
| Adenosine receptors | A1/2A/2B/3 | Active | Active | Active |
| Transmembrane domain-containing protein | TMIGD3 | Active | - | Active |
| E3 ubiquitin-protein ligase Mdm2 | MDM2 | pIC<sub>50</sub>=7.7 | - | - |
| Latent membrane protein 1 | LMP1 | pAC<sub>50</sub>=6.1 | - | - |
| Receptor-type tyrosine-protein kinase | FLT3 | pIC<sub>50</sub>=6.1 | - | - |
| Nuclear receptor coactivator 3 | NCOA3 | pIC<sub>50</sub>=5.1 | - | - |
| DNA (cytosine-5)-methyltransferase 1 | DNMT1 | - | pIC<sub>50</sub>=6.5 | - |
| Peptidyl-tRNA hydrolase | PTH | - | pK<sub>d</sub>=5.8 | - |
| cAMP-dependent protein kinase type I alpha | PRKAR1A | - | - | pK<sub>i</sub>=8.6 |
| Growth hormone secretagogue receptor 1 | GHSR | - | - | pEC<sub>50</sub>=7.3 |
| Dipeptidyl peptidase 4 | DPP4 | - | - | pIC<sub>50</sub>=7.2 |
| Isoleucyl-tRNA synthetase | IleRS | - | - | pIC<sub>50</sub>=7.1 |
| Membrane lipoprotein TmpC | tmpC | - | - | pK<sub>d</sub>=6.6 |
| High affinity choline transporter 1 | SLC5A7 | - | - | pIC<sub>50</sub>=6.2 |
| Histidine triad nucleotide-binding protein 1 | HINT1 | - | - | pIC<sub>50</sub>=6.0 |
| Adenosine kinase | ADK | - | - | pIC<sub>50</sub>=6.0 |
| Mitogen-activated protein kinase kinase kinase 7 | MAPK14 | - | - | pK<sub>d</sub>=5.4 |
| Sodium/nucleoside cotransporters | SLC28A1/3 | - | - | pIC<sub>50</sub>=5.3 |
| DNA polymerase | POL | - | - | Active |

In addition, cytarabine is also known to bind to four other proteins, namely, E3 ubiquitin-protein ligase Mdm2 (MDM2), latent membrane protein 1 (LMP1), receptor-type tyrosine-protein kinase FLT3 (FLT3), and nuclear receptor coactivator 3 (NCOA3). In turn, azacytidine is known to have affinity for DNA (cytosine-5)-methyltransferase 1 (DNMT1) and peptidyl-tRNA hydrolase (PTH). Interestingly, chloroquine, one of the drugs being currently used against Covid-19, is also known to bind to a methyltransferase [23], the histamine N-methyltransferase (HNMT). Finally, adenosine is known to interact with multiple proteins, with most potent affinities for cAMP-dependent protein kinase type I alpha (PRKAR1A), growth hormone secretagogue receptor 1 (GHSR), dipeptidyl peptidase 4 (DPP4), and isoleucyl-tRNA synthetase (IleRS). Altogether, these three drugs provide a list of 23 targets that could be relevant to any molecule within their chemical neighborhood, such as NHC.

### Chemical analogues

Beyond the three drugs discussed above, there are two structurally close molecules that deserve consideration (Figure 3). One of them is cytidine, the endogenous substrate of several nucleoside transporters [24], such as the sodium/nucleoside cotransporter 1 (SLC28A1), the solute carrier family 28 member 3 (SLC28A3), and the equilibrative nucleoside transporter 1 (SLC29A1). And the other one is N6-hydroxyadenosine (NHA), a known active molecule against tyrosyl-DNA phosphodiesterase 1 (TDP1) [25] and also shown to act as a potent antiviral [26]. Again, further analysis of the chemical neighborhood of NHC identifies 4 additional targets and provides evidence of potent antiviral effects.

**Figure 3.**
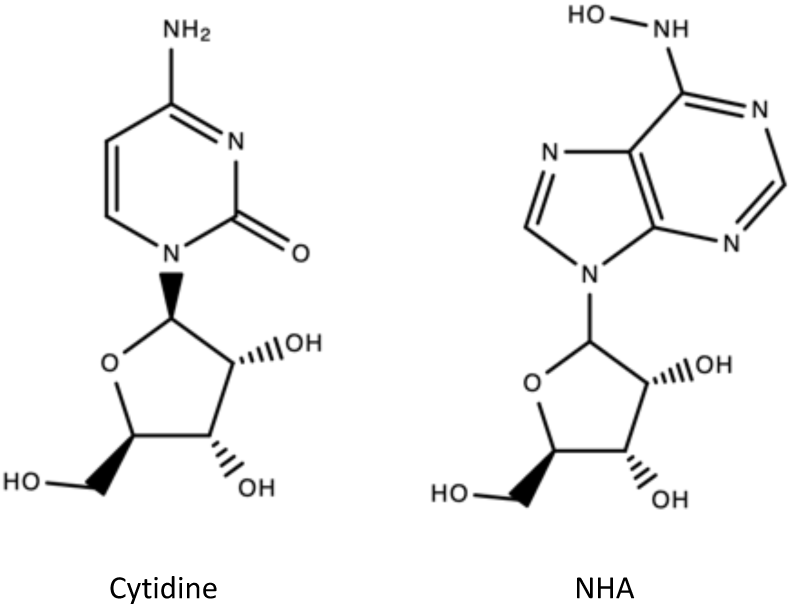
Structures of cytidine and N6-hydroxyadenosine (NHA).

### Predicted metabolites

All metabolic transformations of a given molecule can be also considered part of its chemical neighborhood. Therefore, NHC was processed with CLARITY that predicted two types of metabolic transformations (Figure 4): a hydrolytic deamination, leading to the formation of uridine (with 70% confidence) and a phosphorylation, resulting in the mono/di/tri-phosphate metabolites (with an average confidence of 34%).

**Figure 4.**
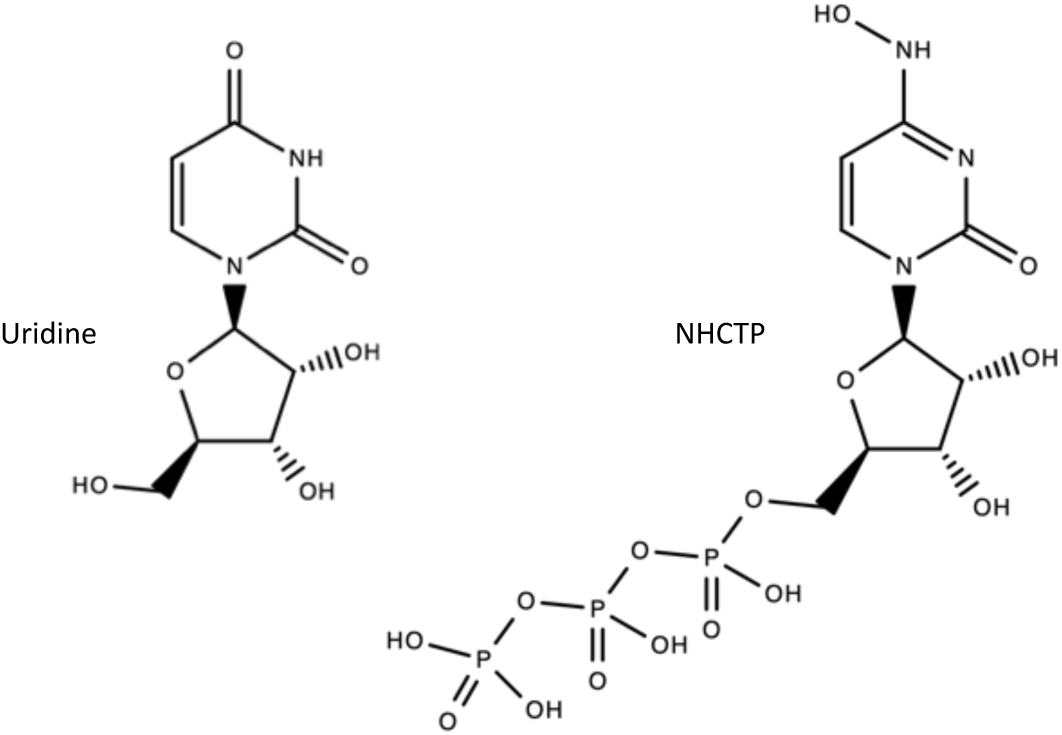
Structures of uridine and the NHC triphosphate metabolite (NHCTP).

Beyond the three nucleoside transporters mentioned above that interact with cytidine, uridine binds to the sodium-dependent nucleobase transporter (SLC23A4), the equilibrative nucleoside transporter 2 (SLC29A2), and the equilibrative nucleoside transporter 3 (SLC29A3) [24]. In addition, uridine is also the result of the catalytic activity of cytidine deaminase (CDA) on its endogenous metabolite, cytidine. The metabolic transformation of NHC into uridine was experimentally confirmed in *Escherichia coli* [27] by measuring the amount of hydroxylamine, a known mutagen [27,28], being released. In turn, uridine can also be transformed into uridine triphosphate that will then bind to several pyrimidinergic receptors, such as P2Y2 and P2Y4 [24].

Finally, the structural similarity of adenosine to NHC will translate into its triphosphate metabolite (ATP) being also in the chemical neighborhood of NHCTP, with all derived consequences in biology. ATP is the endogenous ligands for purinergic (P2X) and pyrimidinergic (P2Y) receptors, it is the substrate or product of numerous enzyme catalytic reactions (such as mevalonate kinase, sphingosine kinases or acetyl-CoA carboxylases) and binds also to several ion channels (such as the transient receptor potential TRPM4 or the inositol and ryanodine receptor channels) [24]. Consequently, all these targets are also likely to interact with NHCTP and they should be added to the list of targets associated with NHC.

### Predicted targets

Following on the recent PHASE initiative [11,12], a list of likely protein targets of NHC was predicted using CLARITY and SEA (Table 2). The highest confidence prediction from CLARITY is for members of the adenosine receptor family, supported by a strong p-value from SEA. This result is very much in line with the only common targets observed within the drug neighborhood (Table 1). In contrast, NHC is most strongly predicted by SEA to bind to P2Y purinoreceptors, whereas no prediction is obtained from CLARITY. A closer look at all results generated reveals that in fact CLARITY predicts those P2Y purinoreceptors not for NHC but for the NHC triphosphate metabolite generated (Figure 4), in agreement with some of the targets known for the adenosine and uridine triphosphates (*vide supra*).

**Table 2.**
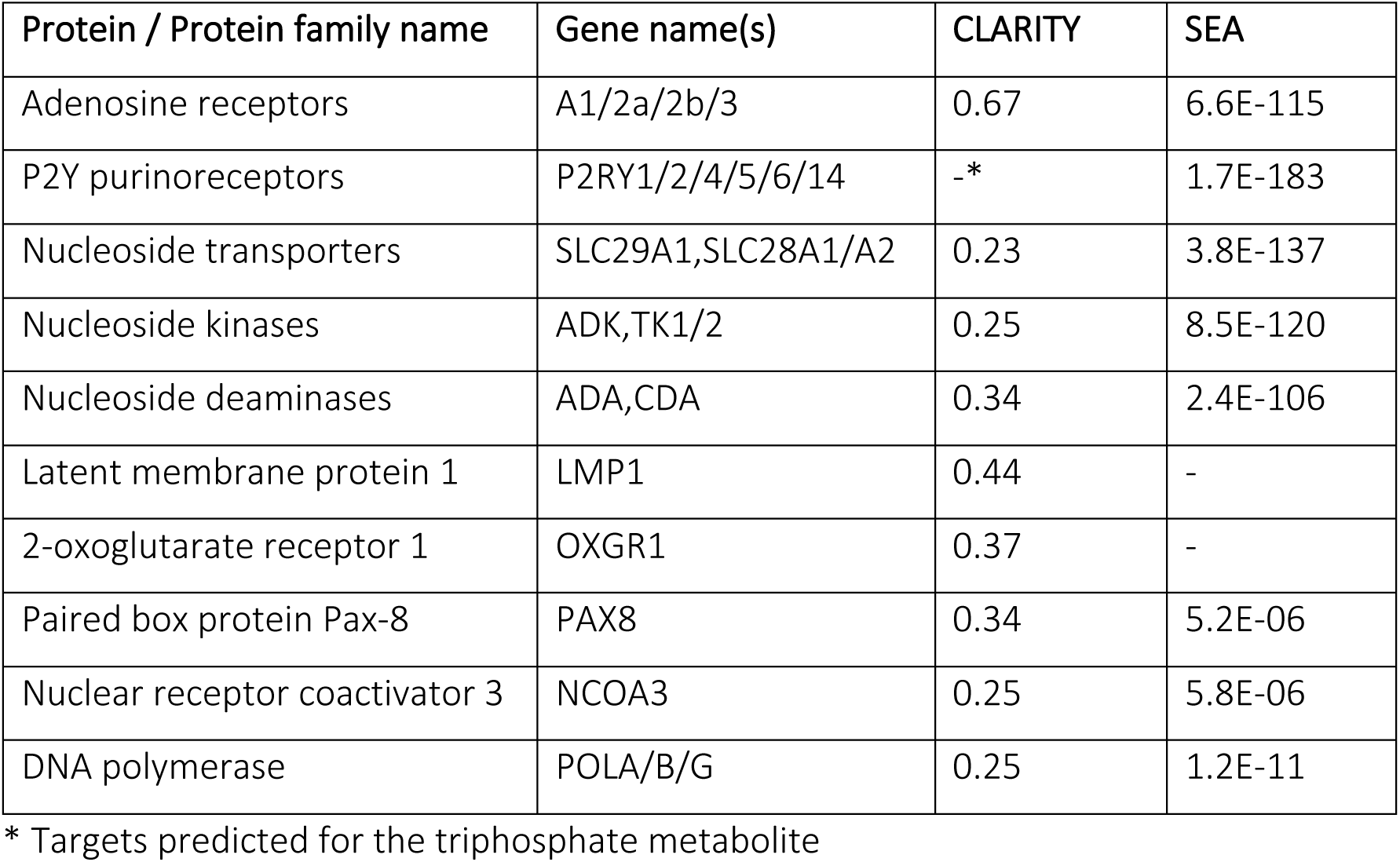
Predicted targets of NHC by CLARITY and SEA. Within a protein family, maximum confidence scores from CLARITY and minimum p-values from SEA are provided.

Among the rest of targets predicted, most of them supported by both platforms, there are some targets known already within the chemical neighborhood of NHC. Those include nucleoside transporters (SLC28 and SLC29) and nucleoside deaminases (such as CDA), known for cytidine, the latent membrane protein 1 (LMP1) and nuclear receptor coactivator 3 (NCOA3), known for cytarabine, and nucleoside kinases (such as ADK) and DNA polymerase, known for adenosine (see Table 1). Predictions highlight also two additional targets not mentioned previously, namely, the 2-oxoglutarate receptor 1 (OXGR1) and the paired box protein Pax-8 (PAX8).

### Relevance to other drugs

All targets identified above may be relevant as well to other antiviral drugs within the same chemical space. This is for example the case of sofosbuvir, that contains the scaffold of uridine, and remdesivir, that has an isosteric scaffold of adenosine. The latter is of particular interest since, even though it was originally designed for treating Ebola [29], remdesivir is one of the drugs being actively pursued in clinical trials against COVID-19 [30]. As it is the case of EIDD-2801, remdesivir is a prodrug of a small molecule nucleoside analogue (GS-441524) that in turn is a precursor to the pharmacologically active triphosphate metabolite (Figure 5) [31].

**Figure 5.**
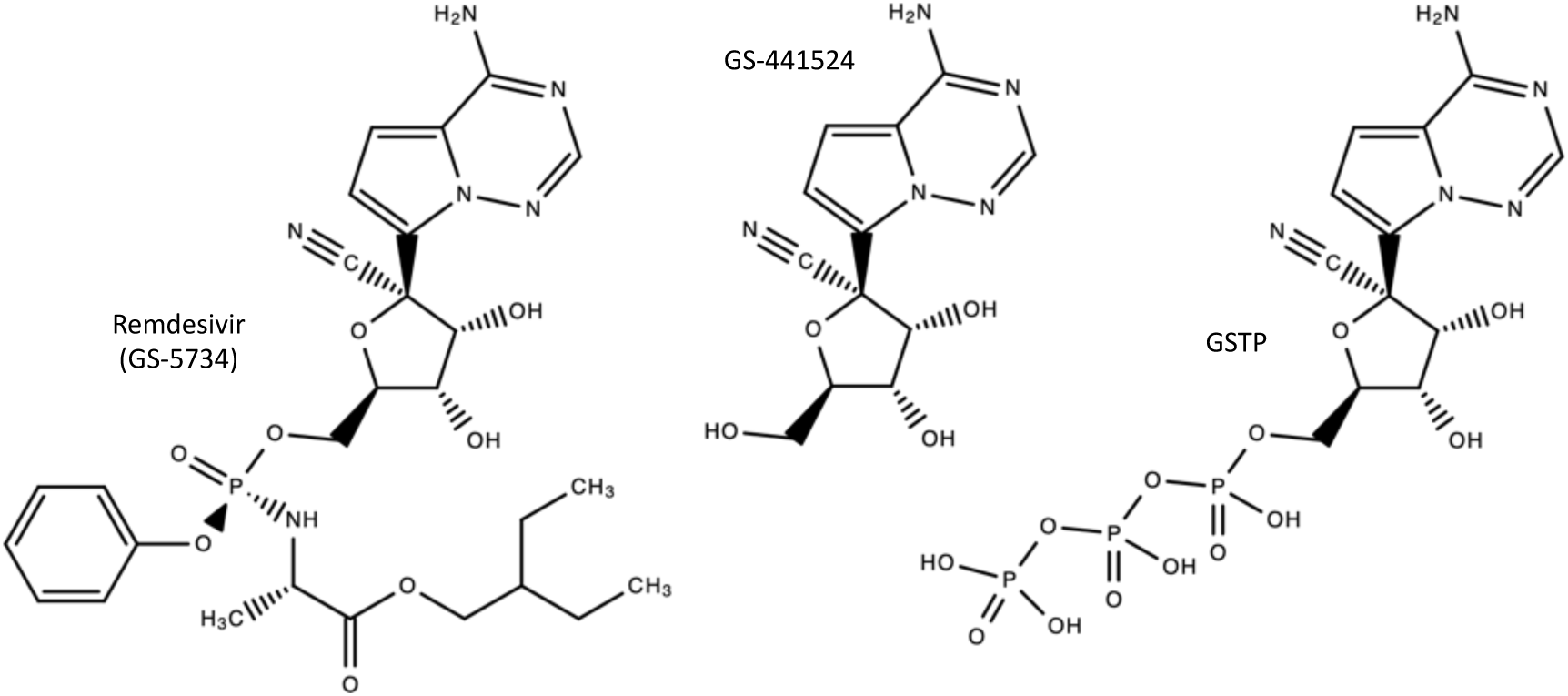
Structures of remdesivir (GS-5734), its nucleoside analogue (GS-441524) and its active triphosphate metabolite (GSTP).

## Conclusions

The current COVID-19 pandemic requires urgent generation of data in multiple facets of the disease, one of them being a priority list of protein targets against which scientists can identify drug repurposing opportunities or design more optimal new chemical entities. In the lack of *in vitro* affinity data across a wide panel of pharmacologically relevant targets one can only revert to *in silico* predictions to identify the likely protein targets for the most promising molecules. Some of these targets, in the eyes of an expert virologist, may hopefully be useful for prioritizing future drug repurposing or discovery campaigns.

## Acknowledgments

I’d like to thank all healthcare professionals for their enormous efforts during this Covid-19 epidemic while we remain safe confined at home trying to contribute with our grain of sand as scientists. This work was supported by a RETOS project from the Spanish Ministerio de Ciencia, Innovación y Universidades (SAF2017-83614-R).

